# Spatial Compartmentalization of TCR Repertoires Between Primary Melanomas and Sentinel Lymph Nodes Reveals Distinct Clonal Architectures and Shared Antigen Recognition

**DOI:** 10.64898/2026.06.05.730356

**Authors:** Simo Kitanovski, Nalini Srinivas, David Schrama, Daniel Hoffmann, Jürgen C. Becker

**Affiliations:** Bioinformatics and Computational Biophysics, Faculty of Biology and Centre for Medical Biotechnology (ZMB), University of Duisburg-Essen, Essen, 45141, Germany; Translational Skin Cancer Research, German Cancer Consortium (DKTK), Essen, Germany; Department of Dermatology, University Medicine Essen, Essen, Germany; German Cancer Research Center (DKFZ), Heidelberg, Germany; Department of Dermatology, University Hospital Würzburg, Würzburg, Germany; Center for Medical Biotechnology, University of Duisburg-Essen, Essen, Germany; Center for Computational Sciences and Simulation, University of Duisburg-Essen, Essen, Germany

**Author notes:** **For correspondence:** (JCB); (SK). Co-last authors (Daniel Hoffmann, Jürgen C. Becker).

## Abstract

Primary tumors and their sentinel lymph nodes are functionally linked sites of anti-tumor immunity, yet how T cell receptor (TCR) repertoires are organized across these compartments remains incompletely understood. We thus performed TCR*β* sequencing on paired primary melanoma tumors and sentinel lymph nodes from 24 treatment-naive patients to quantify TCR repertoire diversity and clonal architecture as markers of antigen-driven selection. Primary tumors exhibited markedly reduced TCR diversity and pronounced clonal dominance compared with matched lymph nodes, consistent with selective expansion of tumor-reactive T cells. Tumor-associated clonotypes displayed significantly longer CDR3 sequences driven by increased non-templated nucleotide insertions, a feature associated with neoantigen recognition, and showed biased TRBV and TRBJ gene usage indicative of CD8^+^ T cell enrichment. Annotation against known melanoma differentiation and cancer testis antigens identified only a small fraction of clonotypes with characterized specificities, and fewer than 10% of these were shared between tumors and lymph nodes, suggesting that most expanded tumor clonotypes recognize patient-specific antigens. To identify shared immune features beyond individual clonotypes, we applied *ClustIRR* to build a joint graph linking clonotypes across repertoires by CDR3 sequence similarity and detected Communities on the Joint graph (CJs) to uncover recurrent TCR sequence motifs. Differential occupancy analysis identified distinct CJs segregating tumors and lymph nodes, including a limited number of recurrent CJs targeting MART-1 epitopes, while the majority were patient-specific. Together, these data define spatially structured TCR repertoire architectures in human melanoma and establish a scalable framework for interrogating tumor-immune interactions.

**eLife Digest:** Melanoma is a skin cancer harboring many UV-associated mutations, making it recognizable to the immune system. T cells attack cancer using specialized molecular structures called T cell receptors (TCRs). They are frequently referred to as specific keys that only fit into the respective locks, i.e. HLA molecules presenting the target antigen. Surgeons often remove both the tumor and the tumor draining lymph node (aka sentinel lymph node). The tumor is where T cells directly fight cancer, while the sentinel lymph node is where new anti-tumor T cells are first activated. How these two sites coordinate immune surveillance is not completely understood.

Here, Kitanovski and colleagues sequenced TCRs from paired tumors and sentinel lymph nodes obtained from melanoma patients. These analyses demonstrated that primary tumors have dramatically reduced TCR diversity, dominated by a few massively expanded T cell clones, whereas sentinel lymph nodes contain broad, balanced repertoires. This reveals intense local selection within the tumor–a small set of tumor-specific T cells expand repeatedly while the lymph node preserves diverse immune reserves.

Examining TCR structure revealed a second key difference. Tumor-infiltrating T cells possess on average slightly longer CDR3 loops–the region contacting antigens–especially among dominant clones. These longer loops arise from random nucleotide insertions. Prior work shows that self-antigen-specific T cells have shorter CDR3s due to thymic constraints, whereas T cells recognizing tumor-specific mutation-derived neoantigens escape these restrictions and have longer CDR3s. Thus, tumor TCRs appear tailored for recognizing patient-specific neoantigens.

The authors could only use TCRs matching known databases for shared melanoma antigens like MART-1 to identify TCR-specific epitopes; most tumor-expanded TCRs could not be annotated because patient-specific neoantigens are not in public databases. This limitation itself is informative– most expanded tumor TCRs likely target private neoantigens.

Grouping TCRs by sequence similarity into “communities” revealed striking patterns: sentinel lymph nodes showed conserved community composition across patients, consistent with responses to common viruses, while tumor repertoires were highly individualized, reflecting each tumor’s unique mutations.

These findings establish how anti-melanoma immunity is spatially organized: sentinel lymph nodes contain diverse T cell landscapes, while in primary tumors melanoma-infiltrating T cells appear to have undergone intense selection for neoantigen recognition. This provides a foundation for using TCR features as biomarkers and designing personalized T cell therapies.

## Introduction

Melanoma derives its strong immunogenicity (***Wang et al., 2019***) from its high UV-induced mutational burden as well as the expression of Cancer-Testis (CTA) and melanocyte differentiation (MDA) antigens (***Alexandrov et al., 2020***). Thus, melanoma is an exceptional model for studying anti-tumor immune responses.

The initial therapeutic management of melanoma typically involves surgical resection of both the primary tumor (PT) and the tumor-draining, or sentinel, lymph node (SLN), rendering these tissues readily accessible for translational investigation. Importantly, PTs and SLNs constitute functionally interconnected (***Lund, 2022***) yet immunologically distinct compartments that together coordinate anti-melanoma immune responses. PTs represent sites of ongoing tumor–immune confrontation, where infiltrating T cells encounter tumor antigens, undergo activation, and expand clonally *in situ*. In contrast, SLNs function as central immunological hubs in which tumor-derived antigens are presented by professional antigen-presenting cells, driving T cell priming, differentiation, and the establishment of memory populations (***Wei et al., 2025***).

Although PTs and SLNs are physiologically linked through lymphatic drainage allowing tumor-derived signals, antigens, and immune modulatory factors to shape SLN immunity and potentially influence metastatic seeding, the extent to which these compartments share or diverge in their T cell receptor (TCR) repertoires remains poorly defined. Recent evidence reveals a critical temporal dimension to this interplay: the primary tumor exerts time-dependent immunosuppressive effects that fundamentally alter the prognostic significance of SLN micrometastatic burden. Specifically, when SLNs are removed immediately or shortly after primary tumor resection, they have enhanced micrometastatic rates, yet paradoxically exhibit reduced immunoactivity characterized by Th17/regulatory T cell predominance; conversely, delayed SLN removal (≥1 week post-PT resection) permits immune reconstitution marked by increased cytotoxic *γδ*T cell frequencies and enhanced CD209+ antigen-presenting cell activity, fundamentally reshaping disease prognostication (***DeTemple et al., 2024***). Systematic, paired analyses of TCR repertoire architecture across matched PT-SLN samples are scarce, leaving a critical gap in our understanding of the spatial and temporal organization of anti-tumor T cell responses.

The TCR repertoire, generated through stochastic V(D)J recombination and subsequently shaped by thymic selection and peripheral antigen exposure, provides a molecular record of immune-tumor interactions. High-throughput sequencing studies have revealed that effective anti-tumor immunity is associated not merely with the presence of individual dominant T cell clones, but with the coordinated expansion and diversification of antigen-specific T cell communities. Such polyclonal responses-characterized by shared antigen specificity, convergent TCR features, and dynamic clonal turnover-have been linked to improved tumor control and responsiveness to immunotherapy. However, how these TCR communities are distributed, maintained, or reshaped between PTs and their corresponding SLNs remains largely unknown.

To address this knowledge gap, we performed deep TCR sequencing on 48 repertoires obtained from paired PT and SLN samples from 24 treatment-naive melanoma patients, integrating these data within a comprehensive multi-modal analytical framework. First, we quantified clonal architecture using normalized rank abundance distributions (NRADs) to assess diversity shifts between compartments. Second, we examined CDR3 structural characteristics-such as length distributions and junctional editing patterns-to identify signatures of antigen-driven selection. Third, we applied Bayesian modeling of TRBV and TRBJ gene usage to uncover compartment-specific biases in V/J recombination. Fourth, we annotated antigen specificities using the VDJdb database to trace clones reactive to melanocyte differentiation antigens (MDAs) and cancer-testis antigens (CTAs). Finally, to transcend the limitations of conventional clonotype-level analyses, we employed *ClustIRR* (***Kitanovski et al., 2026***) to construct a joint graph, a unified network structure where nodes represent TCR clonotypes from all samples and edges connect clonotypes with similar CDR3 sequences regardless of sample origin. Clustering this graph identifies TCR Communities on the Joint graph (CJs) that persist across repertoires, enabling the identification of Differentially occupant CJs (DCJs) reflective of shared antigen recognition that may remain undetected at the individual clonotype level.

Our integrative analysis revealed that PT repertoires exhibit reduced clonal diversity, elongated CDR3 loops, and biased TRBV/J usage-features consistent with intense local antigenic selection. In contrast, SLNs retain broader diversity and preserve signatures of conserved antiviral memory. Community-level analyses further identified recurrent PT-expanded CJs recognizing shared melanoma antigens such as MART-1, mediated through conserved sequence motifs, alongside private CJs likely targeting patient-specific neoantigens. Collectively, this study provides the first quantitative atlas of TCR repertoire dynamics across the PT-SLN axis, offering new insights into the spatial coordination of anti-melanoma immunity and establishing a framework for future biomarker discovery.

## Results

### Biased clonal diversity in TCR repertoires from primary tumors (PTs) and the corresponding sentinel lymph nodes (SLNs)

To assess clonal diversity in TCR repertoires from PT and SLN tissue, we compared their normalized rank abundance distributions (NRADs). This method is a quantitative approach that visualizes the relative frequency of individual T cell clonotypes ranked by their abundance within each tissue. This method effectively captures both the dominance structure (whether a few clonotypes overwhelmingly dominate the repertoire) and the overall diversity (how evenly clonotypes are distributed across the population).

The NRADs form two distinct clusters with different immunological signatures: one comprising PTs (blue curves in Fig. 1) and the other SLNs (orange curves in Fig. 1). Primary tumors exhibited a “top-heavy” distribution with pronounced heads, i.e. the most abundant clonotypes (ranks 1–200) achieved disproportionately high normalized abundance-coupled with relatively weak tails (minimal representation at ranks >500). This pattern indicates strong clonal expansion driven by a small number of dominant TCR clonotypes, consistent with a tumor microenvironment that selectively expands particular T cell populations and thus reducing overall repertoire diversity. Conversely, we see higher entropy of SLN repertoires, or, equivalently, greater T cell clonal diversity–a feature typically associated with functional immune responses capable of recognizing diverse tumor antigens and resisting immune evasion.

**Figure 1.**
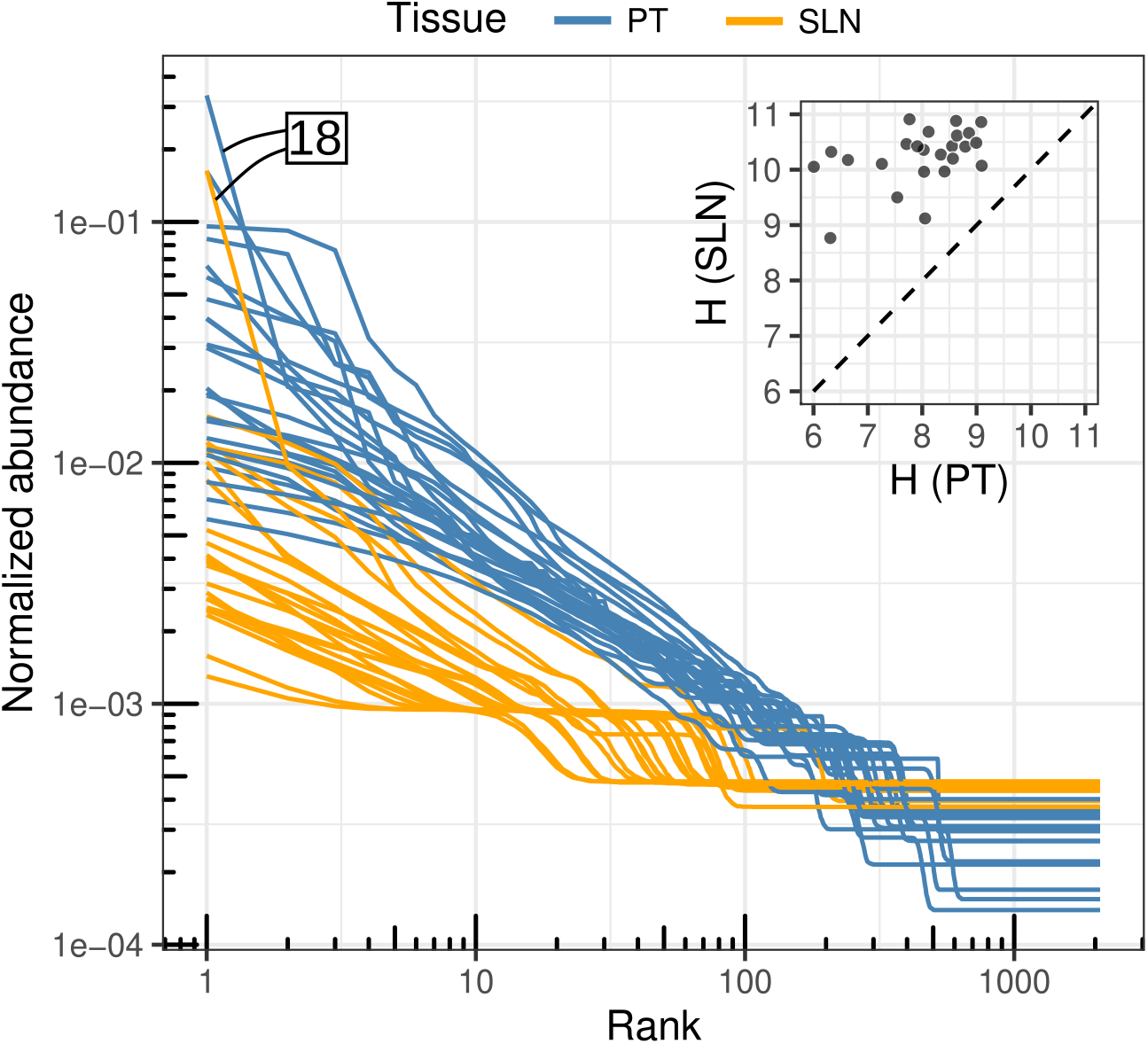
Log-log plot of normalized rank abundance distributions (NRADs) of TCR repertoires from primary tumor (PT; blue lines) and sentinel lymph node (SLN; orange lines) samples. Each line represents an individual patient sample. NRADs reveal distinct clonal architectures: primary tumors display pronounced dominance structures with constrained diversity, while sentinel lymph nodes exhibit flatter distributions reflecting greater clonal heterogeneity. Patient 18 (marked) shows partial concordance at the highest ranks but divergent diversity patterns at lower ranks. An inset scatterplot summarizes Shannon entropy of SLN (y-axis) versus PT (x-axis) for 24 patients, with each point representing a paired PT–SLN sample. The dashed diagonal indicates equal entropy between compartments.

One notable exception emerged in patient 18 for which we observed similar NRADs with strong heads in both tissues. However, while both tissues had similarly pronounced heads (ranks 1–1) indicating comparable clonal dominance at the top of the hierarchy, NRAD curves diverged sharply at ranks > 2, suggesting that both compartments harbored a dominant clonotype, but the SLN had retained substantially greater diversity beyond the most abundant clones. This partial exception highlights intra-individual heterogeneity in PT-SLN immune organization and underscores that clonal dominance patterns may not uniformly predict repertoire architecture across all anatomical compartments in every patient.

To quantify these differences, we calculated the normalized Shannon diversity index (*H*)–a metric that integrates both the number of distinct clonotypes and the evenness of their frequency distribution, providing a single-value summary of repertoire complexity (Fig. 1, inset). Consistent with the NRAD patterns described above, PT samples demonstrated systematically lower diversity compared to SLNs, a finding that confirms and quantifies their substantially reduced TCR complexity.

This divergence in clonal diversity reflects fundamentally distinct immunological functions of these two compartments. Primary tumors exhibit lower TCR diversity because they are sites of active anti-tumor immune selection, where a limited set of tumor-reactive T cell clonotypes undergo iterative expansion in response to persistent tumor-associated antigens. Sentinel lymph nodes, conversely, function as the primary anatomical sites for *de novo* generation and initial clonal expansion of antigen-specific T cells following tumor antigen capture and presentation by dendritic cells. Consequently, SLNs retain higher clonal diversity as they harbor a broader repertoire of nascent tumor-reactive clonotypes with varying specificities, alongside pre-existing bystander T cells that do not recognize tumor antigens but contribute to overall repertoire complexity. Similar patterns have been observed by others in e.g. primary colorectal tumors and their tumor draining lymph nodes (***Matsuda et al., 2019***); or primary lung tumor and non-metastatic regional lymph nodes (***Zhang et al., 2024***).

### Local antigenic pressures shape CDR3 length and junctional diversity in PT versus SLN TCR repertoires

CDR3 length encodes information about T cell receptor genesis, thymic selection stringency, and antigen experience status (***Hou et al., 2019***). CDR3 length distributions therefore serve as a quantifiable descriptor of repertoire composition–distinguishing, for instance, between naïve and memory compartments, CD4 and CD8 subsets, and public versus private clonotypes. Critically, while CDR3 length alone does not predict specificity, systematic differences in length distributions between tissue compartments can reveal selective pressures acting on the T cell repertoire.

Productive TCR repertoires from PTs contained longer CDR3*β* sequences than those from SLNs (Fig. 2A, left panel). Although the difference in mean CDR3*β* length across the entire productive repertoire was modest (Δ_PT−SLN_=0.23, 95% Highest Density Interval (HDI) [0.10, 0.34]), this effect was highly robust (Fig. 2B, top panel). The distinction became substantially more pronounced when analyzing the 100 most expanded clonotypes per sample: here, PT-derived clonotypes were approximately 0.95 nucleotides longer on average than SLN counterparts (Δ_PT−SLN_=0.96, 95% HDI [0.53, 1.37]) (Fig. 2A-B, middle panels), i.e., a fivefold amplification of the signal observed in unselected repertoires. In contrast, non-productive (germline-unconstrained) TCR sequences showed negligible and statistically uncertain length differences between tissues (Δ_PT−SLN_=0.01, 95% HDI [-0.17, 0.19]) (Fig. 2B, bottom panel), demonstrating that the CDR3*β* length disparity is specific to functionally selected repertoires.

**Figure 2.**
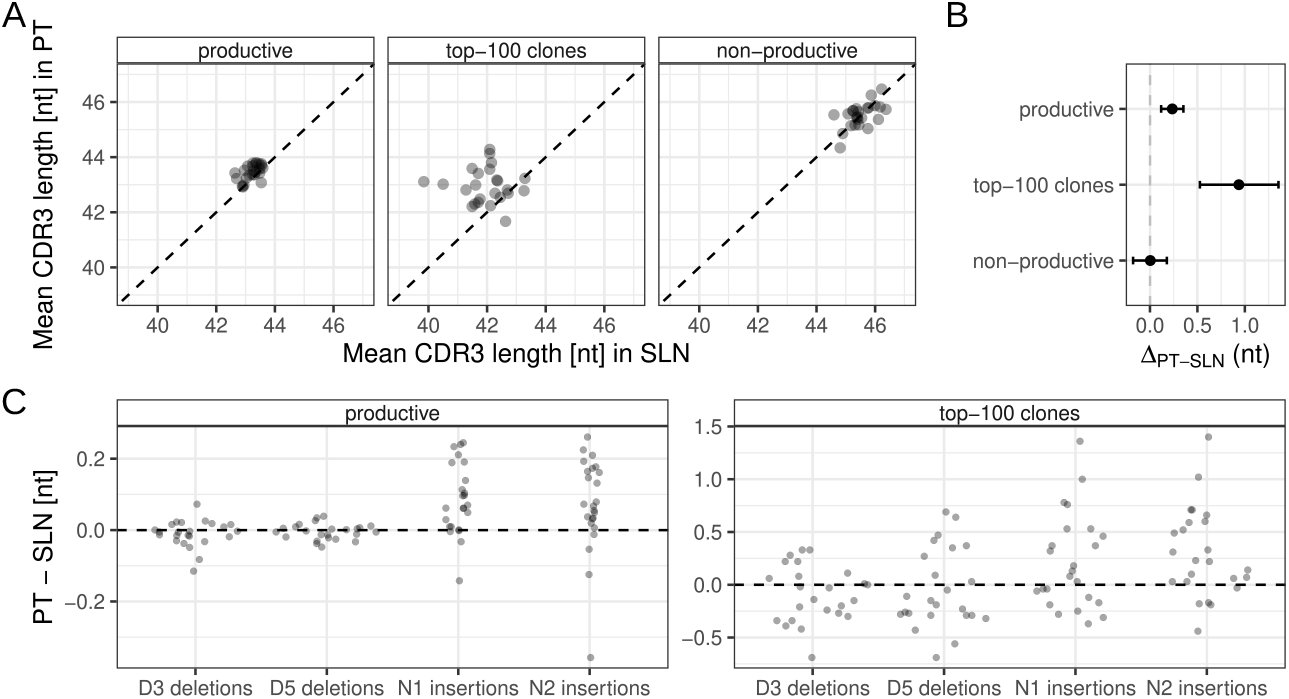
CDR3*β* sequence length and junctional editing differences between PT and SLN TCR repertoires. (A) CDR3*β* lengths in paired primary tumor (PT; y-axis) and sentinel lymph node (SLN; x-axis) TCR repertoires. Panels show CDR3*β* lengths for productive clonotypes (left), the 100 most expanded clonotypes (middle), and non-productive clonotypes (right). Each dot represents a paired PT-SLN sample. The diagonal line (gray) indicates equal lengths. (B) Mean difference (dot) and 95% HDI (black error bars) in CDR3*β* length between tissues (PT vs. SLN) for productive clonotypes (top), the 100 most expanded clonotypes (middle), and non-productive clonotypes (bottom). (C) PT-SLN differences in non-templated nucleotide insertions (N1, N2) and deletions at the 5’ (D5) and 3’ (D3) ends per sample. Panels show differences for productive clonotypes (left) and the 100 most expanded clonotypes (right). Positive values indicate higher insertions/deletions in PTs.

To mechanistically account for longer CDR3*β* sequences in PTs, we quantified non-templated nucleotide insertions (N1 and N2) and junctional deletions flanking the CDR3*β* region (Fig. 2C, left panel). Primary tumors consistently exhibited elevated frequencies of both N1 and N2 insertions relative to SLNs, a pattern that likely explains their longer CDR3*β* loops. By contrast, deletion frequencies at the 5’ and 3’ ends were comparable between tissues. The differential insertion burden was more pronounced among the 100 most expanded clonotypes (Fig. 2C, right panel), suggesting that the selective advantage of longer, N-insertion-rich CDR3*β*s, is particularly pronounced within the dominant tumor-reactive compartment.

The preferential expansion of longer CDR3*β* sequences in PT repertoire especially among dominant clonotypes points to distinct immunological conditions in the tumor microenvironment. We propose three complementary mechanisms: (i) neoantigen-driven selection, whereby longer CDR3*β* loops provide structural flexibility for recognizing diverse, mutated peptide-MHC complexes with variable geometry; (ii) chronic antigen exposure enriching tumor-reactive T cells that characteristically bear elevated N-insertion frequencies and longer junctional regions; and (iii) tissue compartmentalization bias, as PTs are disproportionately populated by tissue-resident effector memory T cells (subject to prolonged local antigen stimulation), whereas SLNs contain predominantly naive or central memory T cells with shorter, less somatically modified CDR3*β* regions. These mechanisms are not mutually exclusive and could operate synergistically to shape PT CDR3*β* length distributions through local antigenic pressure.

This pattern reflects contrasting selection pressures on self-antigen-specific versus neoantigen-specific T cells. Self-antigens impose stringent thymic and peripheral selection constraints, limiting the structural diversity that longer CDR3*β* sequences afford–hence self-reactive tumor infiltrating lymphocytes (TILs) (such as MART-1-specific cells) are enriched for shorter, more germline-proximal CDR3*β* sequences (***Madi et al., 2014***). Conversely, neoantigen-reactive TILs frequently exhibit longer CDR3*β* regions with increased N1/N2 insertions, enabling broader peptide-MHC recognition space and reflecting relaxed selection constraints for T cells that recognize truly novel epitopes (***Hou et al., 2019***). This dichotomy underscores that while CDR3*β* length is shaped by tissue-level antigenic pressures, antigen specificity remains the primary determinant of junctional architecture.

### V/J gene usage reveals CD8^+^ enrichment in primary tumors

Differential TRBV and TRBJ gene segment usage distinguished PT and SLN TCR repertoires. Specifically, 11 TRBV gene segments exhibited strong evidence of differential gene usage (DGU) with posterior probability *π* >= 0.99. Six of these (04-01, 05-05, 06-04, 07-09, 09-01, 15-01) had significantly increased usage in PT relative to SLN, whereas five others (05-01, 06-01, 06-05, 28-01, 30-01) were more used in SLN (Fig. 3A). For TRBJ segments, four genes demonstrated significant DGU: TRBJ02-01 and TRBJ02-07 were markedly enriched in PT, while TRBJ01-01 and TRBJ 01-04 were enriched in SLN (Fig. 3B).

**Figure 3.**
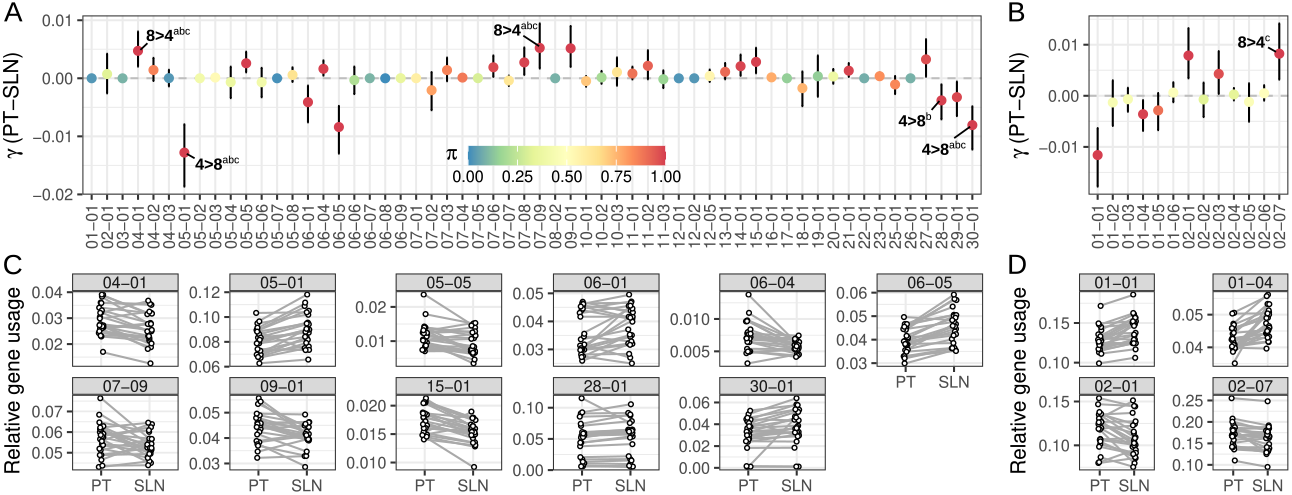
Differential gene usage (DGU) between PT and SLN repertoires. (A) Mean effect size (dot; *γ*) and 95% highest density intervals (HDIs; error bars) for DGU of TRBV gene segments (x-axis) between PT and SLN repertoires. Genes annotated with labels a–c have previously been reported as differentially expressed between CD8^+^ and CD4^+^ T cells (a, ***Miron et al. (2021***); b, ***Li et al. (2024***); c, ***Emerson et al. (2013***)). (B) Mean effect size (dot; *γ*) and 95% HDIs (error bars) for DGU of TRBJ gene segments (x-axis) between PT and SLN repertoires. In panels (A) and (B), dot color indicates the posterior probability of DGU (*π*). (C) Observed relative usage of TRBV gene segments (each panel) in paired PT and SLN repertoires. Dots represent matched PT-SLN pairs. (D) Observed relative usage of TRBJ gene segments (each panel) in paired PT and SLN repertoires. Dots connected by gray lines represent matched PT-SLN pairs.

These tissue-specific TRBV/J biases likely reflect differences in T cell subset composition. SLNs, as secondary lymphoid organs, contain substantial populations of central memory and naive T cells, whereas PTs are enriched for effector/memory clonotypes shaped by local antigen exposure (***Hou et al., 2020***). Moreover, melanoma cells exert immunosuppressive effects on draining lymph nodes, often depleting cytotoxic CD8^+^ T cells while promoting regulatory T cell accumulation (***DeTemple et al., 2024***). To test whether DGU patterns primarily reflect skewed CD8^+^ versus CD4^+^ T cell subset distributions, we compared TRBV and TRBJ usage between human CD8^+^ and CD4^+^ TCR repertoires from publicly available datasets. PT-enriched segments with large positive log-fold-change values (e.g., TRBV07-09, TRBJ02-07) were also significantly upregulated in CD8^+^ versus CD4^+^ repertoires (***Emerson et al., 2013***; ***Li et al., 2024***; ***Miron et al., 2021***) (Fig. 3C and D). Conversely, SLN-preferred segments (e.g., TRBV06-05, TRBJ01-01) showed relative CD4^+^ enrichment. These convergent patterns suggest that the most pronounced differences in TRBV/J usage between PT and SLN likely reflect biases in CD8^+^ versus CD4^+^ TCR repertoires, consistent with the skewed frequencies of these subsets in PT and SLN tissues. However, subset composition alone cannot account for all differences. For example, TRBV09-01 was PT-enriched despite lacking CD8^+^-specific bias, while TRBV06-05 and TRBJ01-01 showed SLN preference without strong CD4^+^ association.

### Public and private antigen-specific clonotypes reveal tumor-localized MDA responses

Clonotype expansion analysis revealed pronounced differences between PT and SLN TCR repertoires, with numerous clonotypes exhibiting greater expansion in PTs relative to SLNs (Fig. 1). To dissect antigen-specific responses, we focused on TCR clonotypes specific for MDAs (MART-1, TRP1/2 and PMEL) and CTAs (NY-ESO-1, MAGE-A3/A4/A6, LAGE-1 SSX2, GAGED2). TCR clonotypes were classified as MDA- or CTA-specific if their CDR3*β* sequences matched entries in VDJdb (Methods). The remaining clonotypes were considered as non-MDA/CTA-specific (MDA-/CTA-).

Across our cohort, we identified 2,063 MDA-specific (43-139 per patient) and 41 CTA-specific (0-4 per patient) TCR clonotypes. MART-1 dominated the MDA response (88%; 1,847 clonotypes), followed by PMEL (10%; 218), with minor contributions from NY-ESO-1 (1.5%; 32) and MAGE-A4/5/6 (0.5%; 9). Relative clonal abundances were comparable between tissues: MDA-specific clonotypes contributed 0.17% of PT and 0.16% of SLN repertoires, while CTAs contributed 0.002% (PT) and 0.003% (SLN). Notably, average clonotype expansion of MDA-specific clonotypes was markedly higher in PTs than SLNs, mirroring patterns observed for non-MDA/CTA-specific clonotypes (Fig. 4A). By contrast, CTA-specific and nonspecific TCR clonotypes showed comparable expansion between tissues (Fig. 4A). Meanwhile, the relative abundance of MDA- and CTA-specific clonotypes and cells were comparable across PTs versus SLNs (Fig. 4B-C). Exceptions included PT repertoires from patients 1, 7, 9, and 20, where MDA-specific cells constituted 0.7-2.6% of total cells, a proportion significantly elevated compared to other PT and SLN samples (Fig. 4C). Patients 6, 7, 9, and 20 additionally exhibited exceptionally high mean clonal expansion of MDA-specific TCR clonotypes in PTs compared to other PT and SLN samples (Fig. 4A).

**Figure 4.**
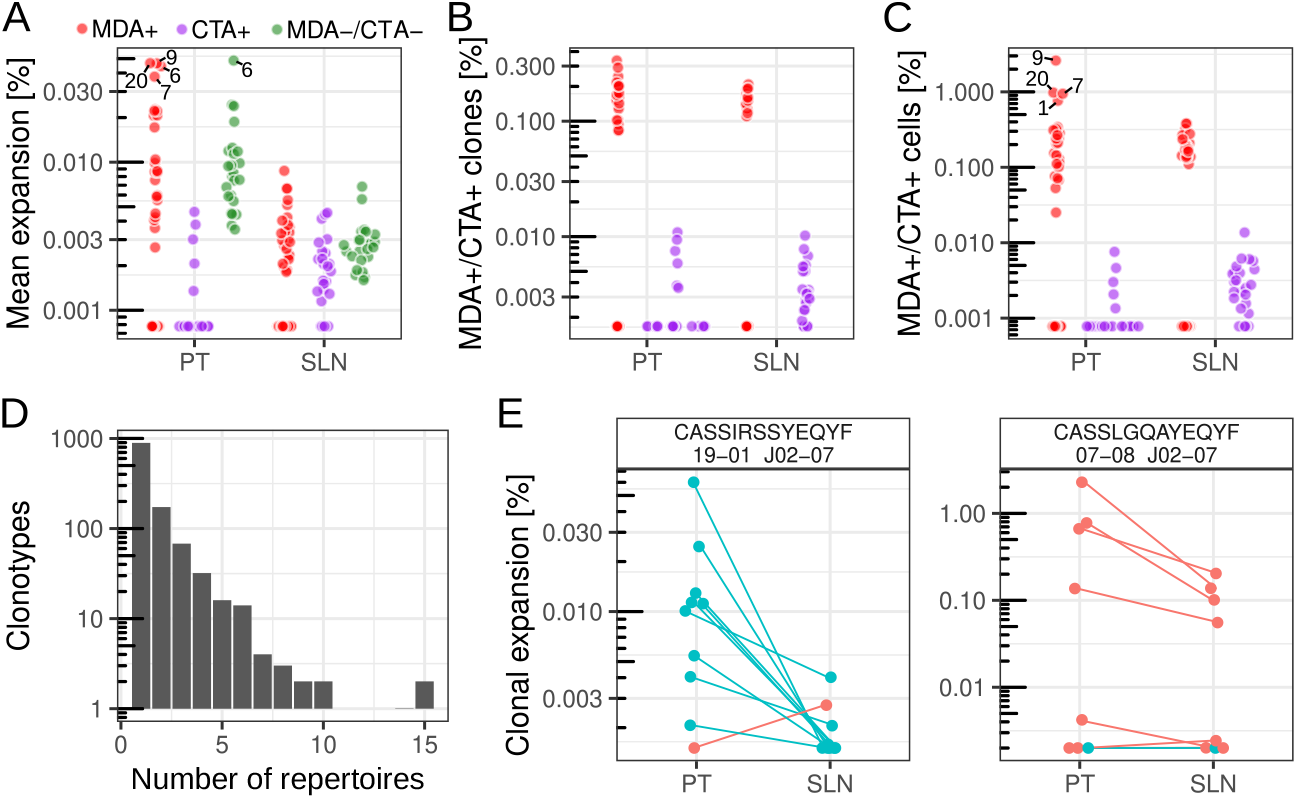
MDA-specific TCR clonotypes. (A) Mean relative expansion of MDA-specific (red), CTA-specific (purple) and non-MDA/CTA-specific (green) clonotypes in PT and SLN repertoires (each dot represents a patient). Patient 20, 9, 6 and 7 are labeled. (B-C) Relative frequency of MDA/CTA-specific clonotypes (panel B) and cells (panel C) in PT and SLN repertoires (dots). Patient 9, 20, 7, and 1 are labeled. (D) Histogram showing the number of repertoires in which each MDA-specific clonotype is detected (x-axis) and the number of clonotypes observed at each level of sharing (y-axis). The majority of clonotypes are private (x = 1), while only two clonotypes are shared across 15 repertoires. (E) Left panel: Expansion (y-axis) of MART-1-specific TCR clonotypes characterized by CDR3*β* sequence CASSIRSSYEQYF, TRBV19-01, and TRBJ02-07, in PT and SLN repertoires (n=10). Green and red dots/segments label HLA-A*02 positive and negative patients, respectively. Right panel: Expansion (y-axis) of MART-1-specific TCR clonotypes characterized by CDR3*β* sequence CASSLGQAYEQYF, TRBV07-08, and TRBJ02-07, in PT and SLN repertoires (n=8). Green and red dots/segments label HLA-B*44 positive and negative patients, respectively.

Among the 2,104 MDA- and CTA-specific clonotypes, only 200 were found in matching repertoires of patients, suggesting a low degree of overlap between matching PTs and SLNs (Fig. 4D). On the other hand, some MDA- and CTA-specific clonotypes were shared across patients, i.e., we found 1,210 unique combinations of CDR3 sequence and TRBV/TRBJ gene segments, including, 893 (74%) private combinations (detected in only one patient), 173 (14%) shared by two patients, and five public shared by 10-15 patients (Fig. 4E).

Only two public clonotypes showed statistically significant PT enrichment relative to SLNs. The first (CDR3: CASSIRSSYEQYF; TRBV19-01; TRBJ02-07) was shared across 10 patients, including 9/11 HLA-A*02^+^ individuals, with robust PT enrichment in all nine HLA-A*02 carriers and incidental SLN detection in one HLA-A*02 negative patient (Fig. 4E, left). VDJdb validation confirmed specificity for the HLA-A*02-restricted MART-1 epitope ELAGIGILTV, highlighting its prominence in anti-melanoma immunity. Meanwhile, TCR*β* repertoire data (***Emerson et al., 2017***) corroborated HLA-A*02 restriction (Supplementary Figure S1). The second public clonotype (CDR3: CASSLGQAYEQYF; TRBV07-08; TRBJ02-07) was shared by 8 patients and validated against the HLA-B*44-restricted MART-1 epitope EEYLKAWTF (Fig. 4E, right). However, expansion was minimal and tissue-comparable, consistent with detection in only 1/4 HLA-B*44^+^ patients.

### CJ composition distinguishes tumor-shaped versus lymphoid-regulated repertoires

Given that the majority of TCR clonotypes are private (>70% in our cohort), a clonotype-centric analysis provides only a limited view of the dynamics of MDA/CTA-specific TCR clonotypes between PTs and SLNs. Notably, TCRs with similar CDR3*β* sequences can recognize related peptide/MHC complexes, a property that is not captured when clonotypes are analyzed individually.

To assess differential occupancy patterns at the level of sequence-related TCRs, we used our recently developed clustering approach *ClustIRR*. This analysis grouped the 1,262,639 TCR clonotypes from 48 repertoires into approximately 636,000 distinct Communities on the Joint graph (CJs) based on shared CDR*β* sequence similarity. Specifically, *ClustIRR* integrated all repertoires into a joint graph where nodes represent clonotypes and edges connect clonotypes with similar CDR3 sequences across samples. CJs were classified as MDA- or CTA-specific if they contained at least one TCR with a corresponding annotation.

To quantify repertoire similarity using CJs, we represented each TCR repertoire as a vector of relative cell abundances (“occupancy”) across all 636,000 CJs and computed pairwise cosine similarity scores, *CS*(*a, b*), where *a* and *b* denote repertoires being compared. This metric captures the directional alignment of CJ occupancy profiles between two repertoires, with scores ranging from 0 (no overlap in CJ composition) to 1 (identical CJ structure and proportions; Fig. 5). SLN repertoires exhibited higher inter-patient similarity scores (mean *CS* = 0.13, minimum *CS* ≈ 0, maximum Number of repertoires *CS* = 0.36) indicating more conserved CJ structures across individuals. This contrasts sharply with PT repertoires, which showed low inter-patient overlap (mean *CS* = 0.01, minimum *CS* ≈ 0, maximum *CS* = 0.17) and thus display “private” CJ architecture reflecting the known heterogeneity of melanoma tumors – distinct antigenic profiles, variable immune infiltration, and patient-specific clonal expansions drive individualized repertoire signatures.

**Figure 5.**
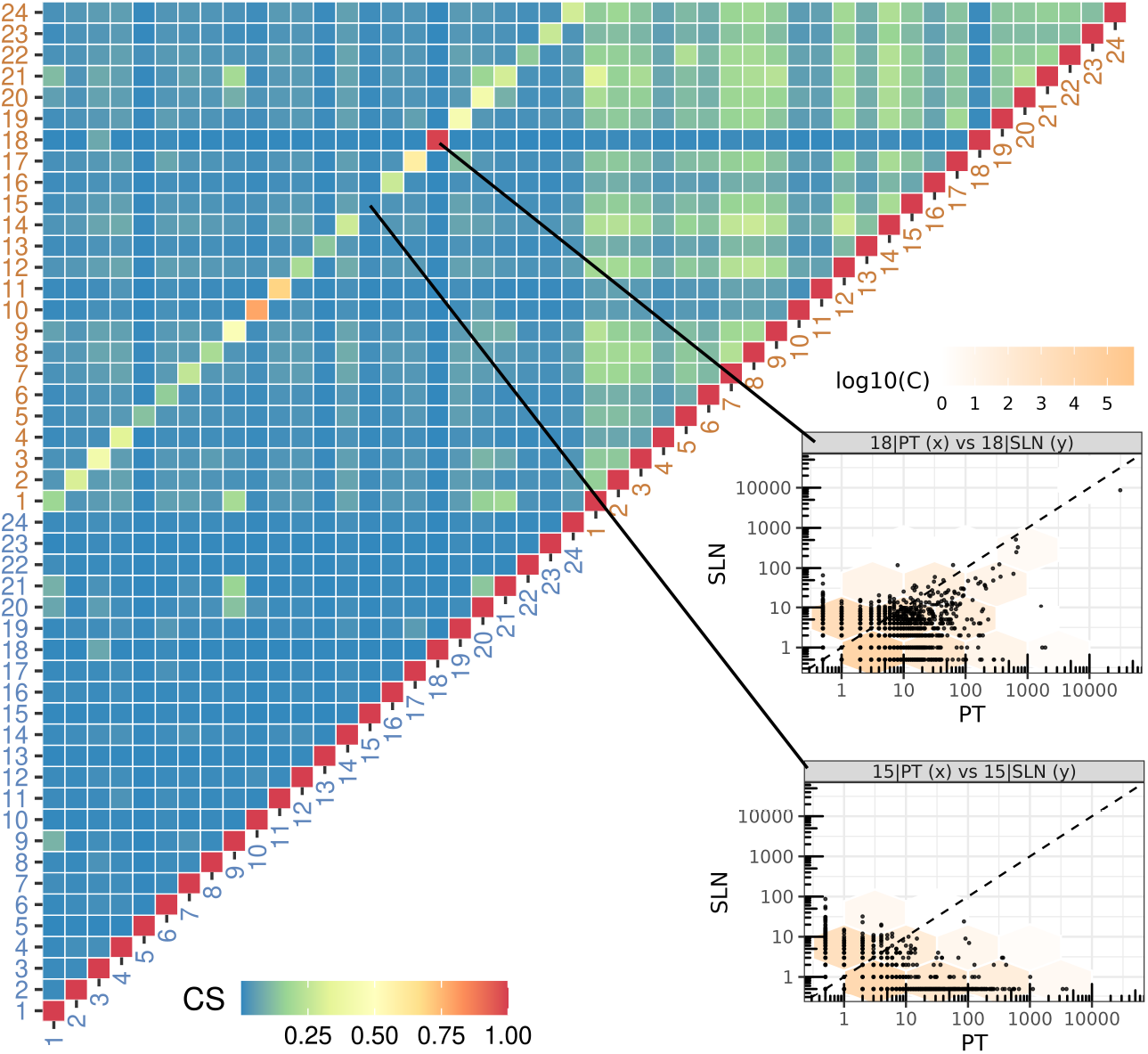
TCR repertoire overlap. Heatmap showing cosine similarity (CS) between pairs of TCR repertoires from primary tumors (PTs; blue labels) and sentinel lymph nodes (SLNs; orange labels), with color-coded tiles representing CS values. Scatterplots showing CJ occupancy between PT and SLN for patients 18 (high CS) and 15 (low CS) are shown in the panels.

The higher SLN-SLN similarity suggests regulation by conserved patient-independent mechanisms, such as homeostatic T cell proliferation and exposure to shared environmental antigens, with minimal influence from tumor-specific pressures. Consistent with this model, similarity of non-related PT-SLN pairs (mean) was lower than for non-related SLN-SLN pairs (mean), emphasizing tumor-localized selection as the dominant driver of PT repertoires structure. However, within-patient PT-SLN overlap (mean *CS* = 0.34) significantly exceeded between-patient PT-SLN overlap (mean *CS* = 0.03) across the cohort (U-test, p-value = ≈ 10^−14^), indicating that patient identity and SLN-PT communication favor repertoire convergence (Supplementary Figure S2). Notably, a subset of patients exhibited exceptional PT-SLN concordance; for example, patient 18 displayed near-perfect similarity (*CS* = 0.99), paralleling their superimposable NRAD curves and shared dominant clonotypes (Fig. 1).

Analysis of differentially occupant CJ To overcome inherent limitations in comparing predominantly private TCR clonotypes between PTs and SLNs, we employed *ClustIRR*’s statistical framework to evaluate T cell occupancy shifts at the CJ level. We defined a Differentially occupant CJ (DCJ) as a patient-specific instance where a global CJ exhibits significant occupancy bias between PT and SLN (*π* ≥ 0.99). Thus, a single global CJ may generate multiple DCJs if it is differentially occupied in multiple patients. Using this approach, we identified 11,030 DCJs corresponding to 10,198 distinct CJs (Fig. 6A). The number of DCJs exceeded the number of unique CJs because recurrent CJs contributed multiple DCJ events (one per patient in which they were differentially occupied). Of these, 8,145 DCJs were expanded in PTs relative to SLNs (positive ϵ), while 2,885 had negative ϵ. The absolute magnitude of ϵ was significantly greater for PT-expanded DCJs than for SLN-expanded counterparts. Notably, only 55 DCJs contained TCR clonotypes specific for MDAs or CTAs. Analysis of DCJ recurrence revealed that the majority were patient-specific: 9,693 DCJs (95%) occurred in only a single patient, whereas 505 DCJs (5%) were shared by ≥two patients (Fig. 6B). Among these shared DCJs, we identified three recurrent DCJ sets expanded across ≥5 PTs but ≤1 SLN (DCJs *r1*–*r3*), and five recurrent DCJ sets expanded across ≥8 SLNs but ≤1 PT (DCJs *r4*–*r8*).

**Figure 6.**
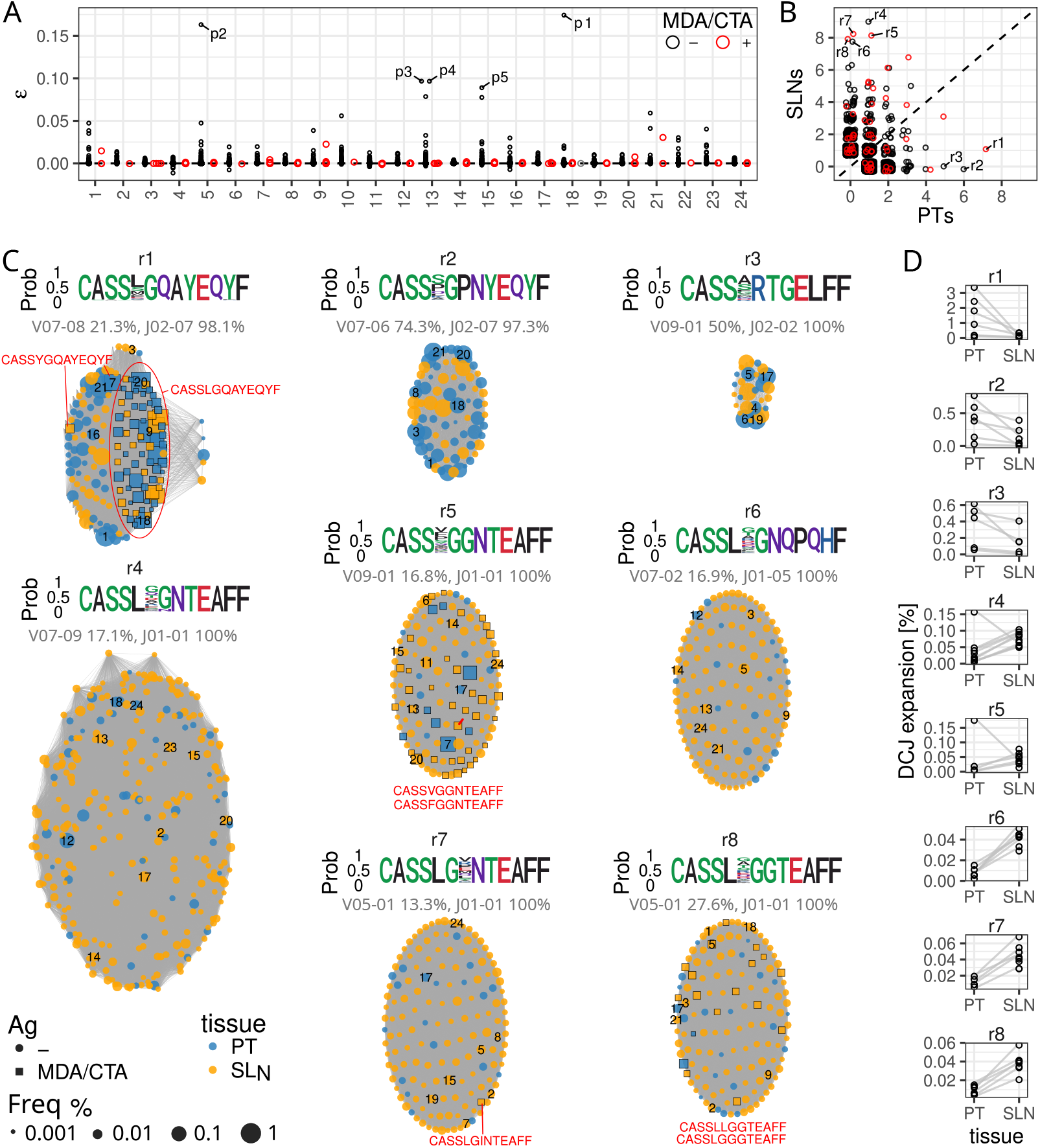
Recurrent Differentially occupant Communities on the Joint graph (DCJs). (A) **Differential occupancy landscape.** Each point represents a DCJ identified between PTs and matched SLNs, plotted by effect size (ϵ) across patients. PT-expanded DCJs (ϵ > 0) appear above the dashed line; SLN-expanded DCJs (ϵ < 0) below. Red points denote DCJs containing at least one MDA- or CTA-specific clonotype; black points indicate non-specific CJs without annotated antigen specificity. Key private DCJs (*p1*-*p5*) are highlighted. (B) **Recurrence of DCJs across patients**. Dot plot showing, for each DCJ, the number of patients exhibiting PT expansion (x-axis) versus SLN expansion (y-axis). Dot colors correspond to panel A. (C) **Network of recurrent DCJs (*r1*-*r8*)**: Nodes represent TCR clonotypes from PTs (blue) and SLNs (orange), with node size proportional to relative clonal expansion in the corresponding repertoire. Gray edges connect clonotypes sharing similar CDR3*β* sequences. MDA/CTA-specific clonotypes are shown as square nodes with black border; whereas non-annotated clonotypes are shown as circles. The most expanded clonotypes per patient are indicated with patient identifiers. Sequence logos depict conserved CDR3*β* motifs within each DCJ set. The most frequently used TRBV and TRBJ gene segments and their usage percentages are shown as gray labels. (D) **Relative DCJ occupancy**. Relative occupancy (%) of DCJs *r1*-*r8* in PTs compared with matched SLNs.

Set *r1* was expanded in the PTs of seven patients and the SLN of one patient (Fig. 6D). It comprised 155 clonotypes, including 72 MART-1-specific clonotypes (≥1 per patient). This set contained a previously identified PT-expanded clonotype (CDR3*β*: CASSLGQAYEQYF, TRBV07-08; TRBJ02-07; Fig. 4E), which targets the HLA-B*44-restricted MART-1 epitope EEYLKAWTF (annotated in VDJdb) and is known to cross-react with Epstein-Barr virus (EBV) antigens. Strikingly, this DCJ featured a prominent cluster of clonotypes sharing identical CDR3*β* sequences and TRBJ02-07 usage but diverse TRBV segments (Fig. 6C). This suggests that a conserved antigen-binding motif can be paired with multiple V-genes while maintaining antigen specificity. Additionally, the set harbored two clonotypes carrying a MART-1-specific variant (CDR3*β*: CASSYGQAYEQYF) targeting the HLA-A*02-restricted MART-1 epitope ALAGIGILTV (VDJdb). Finally, this set included PT-expanded clonotypes with high CDR3 similarity to known MART-1-reactive sequences.

Set *r2*, expanded in the PTs of six patients (Fig. 6D), consisted of 74 clonotypes. PT-expanded clonotypes within this set differed by only one to two amino acids from known PMEL (CDR3*β*: CASSSGPSYEQYF) and MART-1 (CDR3*β*: CASSSGPSDEQYF) variants, suggesting the presence of previously unrecognized MDA-reactive variants. A distinct subgroup within *r2* displayed specificity for cytomegalovirus (CMV; Supplementary Figure S3).

Set *r3*, expanded in the PTs of five patients (Fig. 6D), consisted of 28 clonotypes. While clonotypes within this set did not display specificity to known MDA/CTA antigens, a distinct subgroup demonstrated specificity for EBV (Supplementary Figure S3).

In contrast, sets *r4*–*r8* displayed predominant expansion in SLNs (across seven to nine patients; Fig. 6D), with *r4* and *r5* also expanded in one PT. Sets *r5* and *r8* were enriched with MDA-specific clonotypes (MART-1-specific CDR3*β*s: CASSVGGNTEAFF, CASSFGGNTEAFF, and CASSLGGGTEAFF; PMEL-specific CDR3*β*: CASSLLGGTEAFF), while *r7* contained one MDA-specific clonotype (MART-1-specific; CDR3*β*: CASSLGINTEAFF). All five sets included clusters specific for CMV or EBV (Supplementary Figure S3). Overall, the relative occupancy of SLN-expanded DCJs (*r4*–*r8*) was approximately an order of magnitude lower than that of the PT-expanded sets (*r1*–*r3*), underscoring a marked asymmetry in community-level TCR expansion between PTs and SLNs (Fig. 6D).

The antigen-driven nature of the recurrent DCJs was validated by HLA restriction patterns. Multiple recurrent DCJs that specifically expanded in PTs showed significant enrichment for specific class I HLA alleles. For instance, DCJ set *r2* was strongly associated with HLA types HLA-A*01, HLA-B*08, and HLA-C*07 (Supplementary Figure S4A-C). Independent TCR*β* repertoires corroborated this association, revealing a significantly higher abundance of *r2* CDR3*β* sequences in HLA-A*01+ and HLA-B*08+ individuals compared to HLA-A*01– or HLA-B*08– individuals (Supplementary Figure S4E) (***Emerson et al., 2017***). DCJ set *r1*, enriched for MART-1-reactive CDR3*β*s, showed similar HLA associations, albeit less strongly. The remaining recurrent DCJ sets displayed weaker HLA associations (Supplementary Figure S4). This HLA-restricted enrichment, specific to PT-expanded DCJs, provides compelling evidence that the observed PT/SLN asymmetry reflects antigen-specific T cell responses driven by tumor-associated antigens.

### Patient-specific DCJs reflect tumor-driven antigen recognition

Among DCJs unique to individual patients (private DCJs), we observed pronounced and selective expansion within PTs. These represent global sequence communities that underwent differential expansion in only a single patient, likely reflecting private neoantigen recognition. This was exemplified by DCJs *p1*-*p5* (Fig. 7A), each displaying a relative PT occupancy exceeding 7% (ϵ > 0.7; Fig. 6A). None of these DCJs showed specificity for MDAs, CTAs or common pathogens such as CMV or EBV (Supplementary Figure S5). Intriguingly, however, CDR3*β* sequences within DCJ *p3* exhibited sequence similarity to VDJdb-annotated NY-ESO-1-specific (CDR3*β*: CASSFEGGTGELFF) CDR3*β*, respectively, suggesting recognition of related antigenic motif.

**Figure 7.**
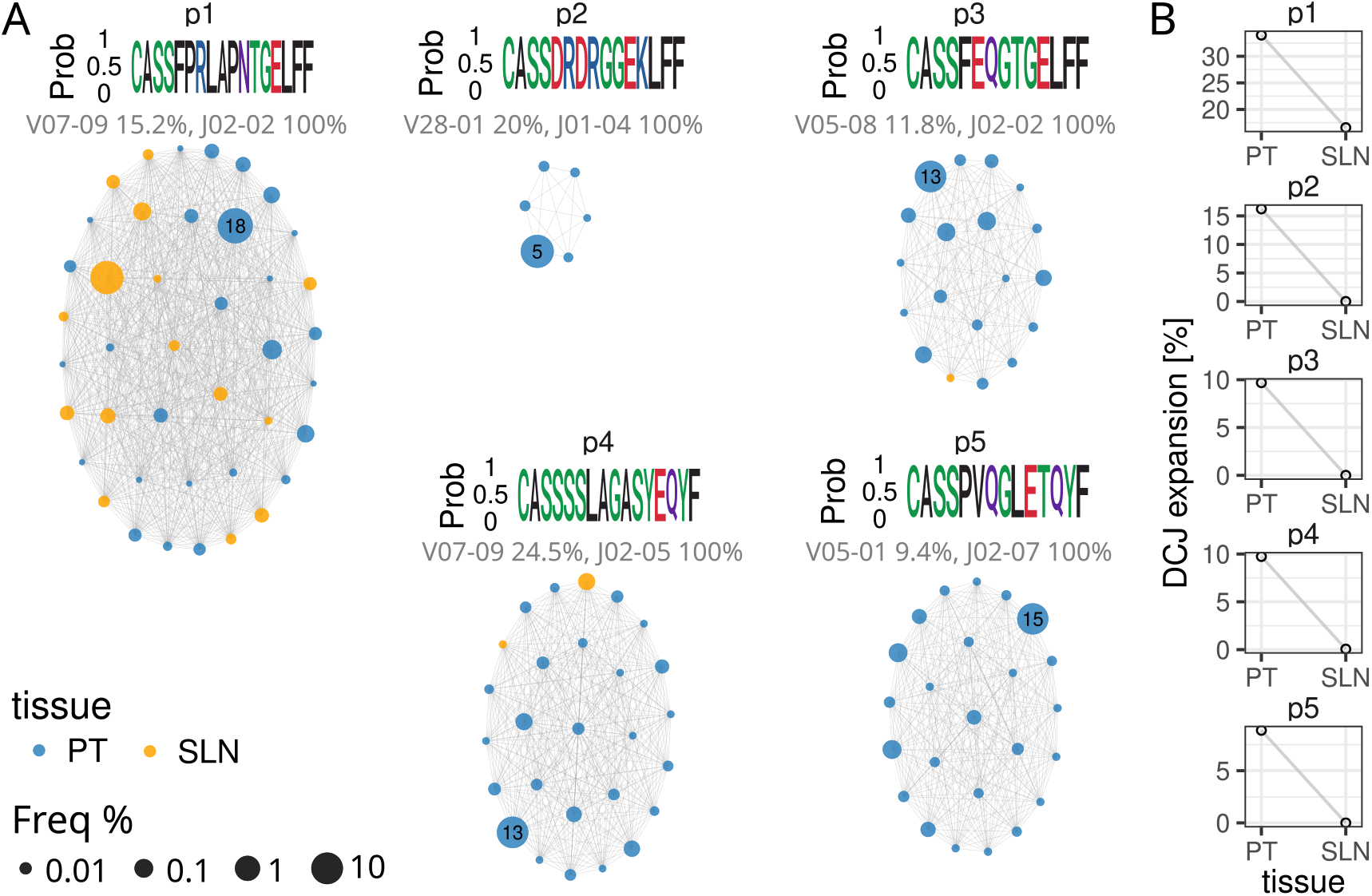
Patient-specific Differentially occupant Communities on the Joint graph (DCJs). (A) **Network of DCJs (*p1*-*p5*)**: Networks depict patient-specific DCJs, visualized using the same conventions as in Fig. 6. Nodes represent individual TCR clonotypes detected in PT (blue) or SLN (orange) with node size proportional to clonal expansion in the corresponding repertoire. Grey edges connect clonotypes with similar CDR3*β* sequences. Black lines highlight the most expanded clonotypes per patient, with corresponding CDR3*β* sequences and patient identifiers indicated. Sequence logos summarize conserved CDR3*β* motifs within each DCJ. The most frequently used TRBV and TRBJ gene segments and their usage percentages are shown as gray labels. (B) **Relative occupancy of *private* DCJs.** Relative occupancy (%) of DCJs *p1*-*p5* in PTs compared with matched SLNs.

Given the absence of active infections in these otherwise healthy melanoma patients, the marked enrichment of these CJs in PTs is unlikely to reflect antiviral responses and instead points to tumor-driven T cell expansion. Our community-level analysis confirms that virus-specific bystander T cells, despite lacking cognate antigen in tumors, undergo substantial clonal expansion and achieve tissue residency through inflammatory recruitment mechanisms (***Rosato et al., 2019***; ***Duhen et al., 2018***). This phenomenon suggests that the tumor microenvironment creates permissive conditions for bystander differentiation, enabling non-tumor-specific memory cells to acquire tumor-infiltrating phenotypes and potentially contribute to anti-tumor immunity through cytokine-mediated activation pathways.

Collectively, our community-level analysis delineates how melanoma heterogeneity shapes distinct patterns of antigen recognition. Recurrent PT-expanded DCJs (*r1*-*r3*) reflect responses to shared self-antigens such as MART-1, that are commonly overexpressed in melanoma. In contrast, patient-specific DCJs with strong PT enrichment (e.g., *p1*-*p5*) likely target neoantigens, as their private occurrence aligns with individual mutational landscapes. Finally, SLN-enriched DCJs (*r4*-*r8*) predominantly represent compartmentalized antiviral memory, which also appear to be instrumental in immunological defense against melanoma. This continuum–from shared self-antigen recognition to private neoantigen responses as well as viral-specific bystanders–captures the antigenic hierarchy of melanoma and underscores how tumor heterogeneity drives personalized TCR repertoires directed against tumor-specific targets.

## Discussion

This study is the first quantitative paired analysis of TCR repertoire architecture across the PT– SLN axis, revealing that these compartments exhibit strikingly distinct clonal organizations despite their functional interconnection. Primary tumors display markedly lower clonal diversity with pronounced dominance structures (top-heavy normalized rank abundance distributions), while SLN maintain substantially broader TCR diversity. This architectural divergence reflects fundamental immunological distinctions: PTs represent sites of active, ongoing clonal expansion driven by persistent tumor-associated antigens, whereas SLNs function as primary induction sites where diverse populations of naïve and central memory T cells undergo initial antigen presentation and priming (***Esterházy et al., 2019***). Similar patterns have been observed in colorectal and lung tumors, suggesting a generalizable principle of tumor-immune compartmentalization.

PT-derived clonotypes exhibited systematically longer CDR3 sequences compared to SLNs, with the effect size amplifying fivefold when analyzing the 100 most expanded clonotypes. This lengthening is driven by elevated non-templated nucleotide insertions (N1/N2) at the junctional regions, without corresponding deletion differences. This junctional editing signature likely reflects two complementary mechanisms: (i) neoantigen-driven selection, wherein longer CDR3 loops provide structural flexibility to recognize the diverse geometry of patient-specific mutated peptide–MHC complexes that escape the stringent selection constraints imposed on self-antigens (***Cagiada et al., 2025***); and (ii) chronic antigen exposure, enriching terminally differentiated effector memory T cells that characteristically have more often N-insertions from repeated antigenic stimulation. Notably, self-antigen-specific T cells targeting melanocyte differentiation antigens had shorter CDR3s, consistent with stringent positive and negative selection during thymic education. Thus, our observations illustrate the principle that, while tissue-level antigenic pressures shape junctional diversity at the population level, antigen specificity remains the primary determinant of individual TCR architecture.

Differential TRBV and TRBJ gene usage distinguished PT and SLN repertoires, with six TRBV segments significantly enriched in PTs and five in SLNs. Cross-validation against publicly available human CD8^+^ and CD4^+^ TCR datasets revealed that PT-enriched segments showed strong CD8^+^ bias, while SLN-preferred segments showed relative CD4^+^ association (***Fu et al., 2019***). These convergent patterns indicate that skewed CD8^+^ versus CD4^+^ T cell subset distribution–consistent with melanoma’s known capacity to selectively deplete cytotoxic T cells in draining lymph nodes (***Chemla et al., 2026***)–is a primary driver of differential V/J usage.

Only 200 of the 2,104 identified melanoma-differentiation antigen- and cancer-testis antigen-specific clonotypes were shared between matched primary tumor and sentinel lymph node pairs, highlighting a fundamental paradox: although shared self-antigens are broadly overexpressed in melanoma, anti-tumor T cell responses are dominated by patient-specific repertoires. Analysis of two validated public TCR clonotypes recognizing MART-1 epitopes illustrates this principle (***De-Witt III et al., 2018***). One clonotype, restricted by HLA-A02, was detected in the majority of HLA-A02 positive patients and showed consistent enrichment within primary tumors, indicating a conserved and recurrent component of anti-MART-1 immunity. In contrast, a second clonotype, restricted by HLA-B44, exhibited limited expansion and was detected in only a minority of HLA-B44 positive individuals. This divergence suggests that immunodominance hierarchies are not determined solely by antigen expression or HLA restriction, but are further shaped by factors such as epitope processing efficiency, TCR–peptide–MHC binding properties, and stochastic events during early T cell priming.

The rarity of shared antigen-specific clonotypes between compartments indicates that while both generate tumor-reactive responses, they do so through largely independent clonal selections. This likely reflects divergent antigen sampling–tumors present the complete repertoire of mutated and amplified self-antigens, while SLNs receive filtered tumor lysate–or temporal dynamics wherein early SLN priming generates initial clonal seeds subsequently outcompeted in tumors by higher-affinity neoantigen-reactive populations (***Lee et al., 2020***).

The limitation that a majority of TCRs are private (here: >70%), motivated the use of *ClustIRR*, which clusters TCRs by CDR3 similarity into 636,000 distinct CJs. Remarkably, CJs revealed fundamental differences in inter-individual organization: SLN repertoires were relatively similar between patients, indicating conserved homeostatic regulation by patient-independent mechanisms. In contrast, PT repertoires were relatively dissimilar between patients, reflecting the known heterogeneity of melanoma. Critically, within-patient PT-SLN similarity significantly exceeded between-patient cross-compartmental similarity, demonstrating that patient identity and SLN-PT communication limit divergence of SLN and PT repertoires despite compartmental specialization.

Numerous CJs were differentially occupant between PTs and SLNs, with the majority preferentially expanded in tumors. Only a very small fraction of these CJs could be linked to previously annotated antigen specificities, indicating that most expanded tumor-associated CJs likely recognize antigens that are so far uncharacterized, consistent with patient-specific neoantigens. In contrast, lymph node-expanded CJs more often reflect responses to ubiquitous environmental or latent viral antigens. Despite this overall diversity, a limited number of recurrent CJ patterns were observed across multiple patients. Prominent among these were tumor-expanded CJs enriched for MART-1 reactive TCRs, which comprised diverse clonotypes sharing highly similar sequence motifs. In some cases, identical antigen-binding motifs were paired with multiple TRBV genes, illustrating how convergent immune recognition can arise through distinct recombination pathways. Closely related variants differing by only one or two amino acids were also detected, suggesting the presence of previously unrecognized MART-1 reactive TCRs. In contrast, CJs preferentially expanded in SLNs were dominated by antiviral specificities, consistent with compartmentalized memory responses and supporting a model in which bystander memory T cells are preferentially recruited or retained in lymphoid tissues rather than undergoing selective expansion within tumors.

These findings establish TCR repertoire architecture as a biomarker layer for melanoma prognosis and immunotherapy response (***Chiffelle et al., 2025***). The robust identification of two public MART-1-specific clonotypes provides consensus targets for adoptive T cell therapy and vaccine design. Simultaneously, the predominance of patient-specific neoantigen-reactive DCJs underscores why personalized neoantigen-based approaches may achieve superior outcomes compared to one-size-fits-all strategies. CDR3 length and N-insertion frequency could serve as accessible biomarkers of neoantigen engagement.

Key limitations include restriction to treatment-naive, surgically-resected specimens (***Yuan et al., 2025***); lack of functional validation for uncharacterized CJs; and exclusive focus on TCR*β* repertoires (***Seo and Choi, 2025***). Integration with single-cell RNA-seq would correlate TCR clonotypes with transcriptional state and functional capacity (***Wang et al., 2025***). Despite these limitations, this study provides the first quantitative, integrative analysis of TCR dynamics across the PT-SLN axis, revealing that these compartments represent functionally distinct yet interconnected immunological domains. Community-level analysis transcends clonotype limitations, identifying both recurrent public CJs targeting shared self-antigens and patient-specific neoantigen-reactive populations that constitute the antigenic hierarchy of melanoma. These findings establish a framework for next-generation biomarker discovery and personalized immunotherapy design in melanoma and other cancers.

## Methods and Materials

### TCR*β* analysis by high-throughput sequencing

Genomic DNA (gDNA) was extracted from each patient from primary tumors and sentinel lymph nodes. Following the ImmunoSeq (Adaptive Biotechnologies, Seattle, WA) protocol, the CDR3 region of the TCR*β* chain was amplified and sequenced. In a first polymerase chain reaction (PCR), highly optimized multiplex PCR primers were used to amplify the CDR3 region resulting from a V, D, and J gene rearrangement. After a second PCR, universal adaptor sequences and DNA barcodes allowed identification and high-throughput sequencing on an Illumina MiSeq using the MiSeq ReagentKit v3 150-cycle (Illumina, San Diego, CA). Subsequently, the ImmunoSeq Analyzer (Adaptive Biotechnologies, Seattle, WA) software was applied for quality checking and clustering to eliminate PCR and sequencing errors from downstream analysis.

The resulting TCR repertoires contained for each clonotype: CDR3*β* amino acid sequence, TRBV and TRBJ gene segment annotations, and template count (quantifying clonal expansion). Only productive rearrangements (in-frame sequences without stop codons) were retained for downstream analyses, which included: (1) clonotype diversity assessment, (2) CDR3*β* sequence length analysis, (3) differential TRBV/TRBJ gene usage quantification, and (4) TCR community clustering and differential occupancy analysis.

### TCR repertoire diversity quantification

To quantify the clonotype diversity of TCR repertoires we employed two strategies. First, we characterized the TCR clonotype diversity using Normalized Rank Abundance Distributions (NRADs) (***Saeedghalati et al., 2017***). Normalization of the rank-abundance-distribution (RAD) of T cell clonality was done by MaxRank normalization as implemented in R-package *RADanalysis* (version 0.5.5). Here, the MaxRank was set to the minimum dimension (2,079) of rank abundance vectors for all TCR repertoires. Normalized RADs (NRADs) were computed by 500-fold averaging, and were visualized with R-package *ggplot2* (version 3.5.1). Second, we estimated the Shannon entropy (*H*):

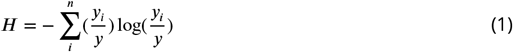

where *y*_*i*_ is the number of cells in clonotype *i*; *y* as the depth of the repertoire with 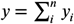.

### Differential TRBV/TRBJ gene usage analysis

Gene segment usage was defined as the count of clonotypes annotated with a specific TRBV or TRBJ segment. Differential gene usage (DGU) between paired PT-SLN samples was quantified using IgGeneUsage (v1.17.27) (***Kitanovski and Hoffmann, 2020***), which employs a Bayesian hierarchical model.

The posterior of the IgGeneUsage model was inferred with rstan (***Carpenter et al., 2017***), using the No-U-Turn sampler in a Markov chain Monte Carlo (MCMC) simulation with 5 chains of 6,000 iterations each, including 2,000 warmups. We assessed the validity of the model with posterior predictive checks (***Gelman et al., 2013***) (Supplementary Figure S6). Furthermore, we inspected the potential scale reduction factor (R-hat), the effective number of samples in the bulk and tails of the posteriors (ESS-bulk and ESS-tail) and information about divergences during the MCMC sampling to check for a successful convergence.

IgGeneUsage outputs mean and 95% highest density interval of the DGU effect size *γ*, and the posterior probability *π* of DGU. TRBV/TRBJ segments with *π* ≥ 0.99 were considered differentially used, where *π* ≈ 1 indicates strong evidence for DGU, while *π* ≈ 0 suggests negligible evidence (not equivalent to absence of DGU).

### CJ detection and differential occupancy analysis

TCR*β* clonotypes were clustered into CJs using *ClustIRR* (R-package, version 1.11.1) (***Kitanovski et al., 2026***) based on CDR3*β* sequence similarity. The *ClustIRR* pipeline executed four sequential steps described in the following.

#### TCR similarity computation

To identify similar TCRs, *ClustIRR* employs protein Basic Local-Alignment Search Tool (*blastp*) via the R-package *rBLAST* (version 1.4.0). Briefly, we constructed a protein database from all CDR3*β* sequences, and each CDR3*β* sequence was used as a query, retaining only CDR3 sequences matches with ≥ 90% sequence identity to the query. For each matched CDR3 pair we computed an alignment score (*ω*) using BLOSUM62 substitution matrix. Identical or highly similar CDR3 sequence pairs receive large positive *ω* scores, while dissimilar pairs receive low or negative *ω*. Scores were normalized by alignment length 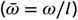.

#### Joint similarity graph construction

*ClustIRR* constructed a graph for each TCR repertoire, representing clonotypes as graph nodes with undirected edges connecting pairs of nodes based on CDR3*β* similarity scores 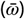. To integrate all repertoires, *ClustIRR* computed similarities between CDR3*β* sequences across different repertoires, creating new edges between nodes from different graphs with 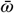 as edge weights. The resulting joint graph (*J* ) contained all 48 TCR repertoires and was structured as an *igraph* object (R-package, version 2.1.1).

#### Community detection

Communities were identified in *J* using the Leiden algorithm (*igraph* implementation) with the Constant Potts Model quality function (resolution parameter=1) over 1,000 optimization iterations. This generated a CJs occupancy matrix *y*^*K*×*R*^ (*K* ≈ 636, 000 CJs; *R* = 48 repertoires) where entries indicate cell counts per CJs-repertoire pair.

#### Differential CJ occupancy analysis

For each patient, paired PT-SLN CJ occupancy vectors were extracted from *y*^*K*×*R*^. Differential occupancy was modeled independently for each patient. CJs with zero cells in both tissues were excluded. Differential occupancy was quantified using a Dirichlet-Multinomial model in *ClustIRR*, which estimates: (1) the effect size (ϵ) for relative occupancy change between PT and SLN (posterior mean, median, and 95% HDI), and (2) the posterior probability (*π*) of differential CJ occupancy. CJs with *π* ≥ 0.99 in a given patient were classified as Differentially occupant CJs (DCJs) for that patient.

The posterior of the *ClustIRR* model was inferred with rstan (***Carpenter et al., 2017***), using the No-U-Turn sampler in a Markov chain Monte Carlo (MCMC) simulation with 4 chains of 1,500 iterations each, including 500 warmups. To test the validity of our model, we performed posterior predictive checks (Supplementary Figure S7). Furthermore, we inspected the potential scale reduction factor (R-hat), the effective number of samples in the bulk and tails of the posteriors (ESS-bulk and ESS-tail) and information about divergences during the MCMC sampling to check for a successful convergence.

### CJ similarity quantification

Cosine similarity (CS) was computed between repertoire pairs using columns from the CJ occupancy matrix *y*^*K*×*R*^. For vectors *A* and *B* representing two repertoires:

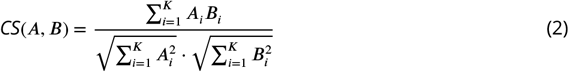

where *A*_*i*_ and *B*_*i*_ are cell frequencies of CJ *i*. Scores range from 0 (no overlap) to 1 (identical composition) given non-negative frequencies.

### Detecting HLA-type enrichment in CJs

To identify CJs enriched for specific HLA types (A/B/C), we compared HLA prevalence within each CJ to its background prevalence across all TCR repertoires using a Bayesian framework. We modeled HLA type usage with a binomial probability mass function: for CJ *j*, the count of clonotypes with HLA type *i* (*y*_*ij*_ ) follows

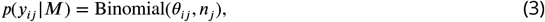

where *θ*_*ij*_ is the HLA type *i* probability in CJ *j* and *n*_*j*_ is the total clonotypes in CJ *j*. Similarly, the global usage of HLA type *i* across all repertoires (*N* clonotypes) is

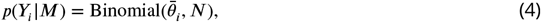

with 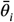 denoting the background probability.

Independent Beta(1,1) priors (uniform distributions) were assigned to *θ*_*ij*_ and 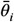. We computed the absolute HLA prevalence difference 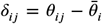. CJs were classified as enriched if *δ*_*ij*_ > 0 and the 95% highest density interval (HDI) for *δ*_*ij*_ lay entirely above zero; depleted if *δ*_*ij*_ < 0 and the 95% HDI lay entirely below zero.

The model was implemented in Stan (***Carpenter et al., 2017***) using the `rstan` package (version 2.32.6). We ran four Markov chain Monte Carlo (MCMC) chains with 2,000 iterations each (1,000 warm-up), employing the No-U-Turn sampler. Convergence was assessed via 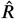 (potential scale reduction factor), effective sample sizes (ESS-bulk and ESS-tail), and divergence diagnostics. Posterior means, medians, and 95% HDIs were extracted as primary outputs, with posterior predictive checks confirming model validity.

### Antigen specificity annotation

Clonotype specificities were annotated via exact CDR3*β* sequence matching against VDJdb (June 2024 download; https://vdjdb.cdr3.net). Only identical matches were considered, with corresponding antigen annotations transferred to matching clonotypes in our dataset. Melanoma differentiation antigens (MDAs) and cancer-testis antigens (CTAs) were specifically flagged for downstream analysis.

### Software and data availability

The TCR*β* repertoire data are available from the corresponding author upon request. The processed data and source code required to reproduce the findings are available at https://github.com/snaketron/ClustIRR_melanoma_paper_data. All figures were generated using *ggplot2* (***Wick-ham, 2016***) and assembled using *patchwork* (***Pedersen, 2025***).

## Acknowledgments

This work was supported by Deutsche Forschungsgemeinschaft (DFG, German Research Foundation) GRK2762 - project number 450917483 (subproject M1) to Daniel Hoffmann. Nalini Srinivas is supported by the Else-Kröner-Fresenius Stiftung (EFKS), University Medicine Essen Medical Scientist Academy (UMESciA). Jürgen C. Becker is supported by the BMFTR DKTK fund ED03. We acknowledge support from the Open Access Publication Fund of the University of Duisburg-Essen.

