## Supplementary for "Spatial Compartmentalization of TCR Repertoires Between Primary Melanomas and Sentinel Lymph Nodes Reveals Distinct Clonal Architectures and Shared Antigen Recognition"

This document includes:
- Supplementary Figures S1-S7
- Supplementary References

### Supplementary Figures

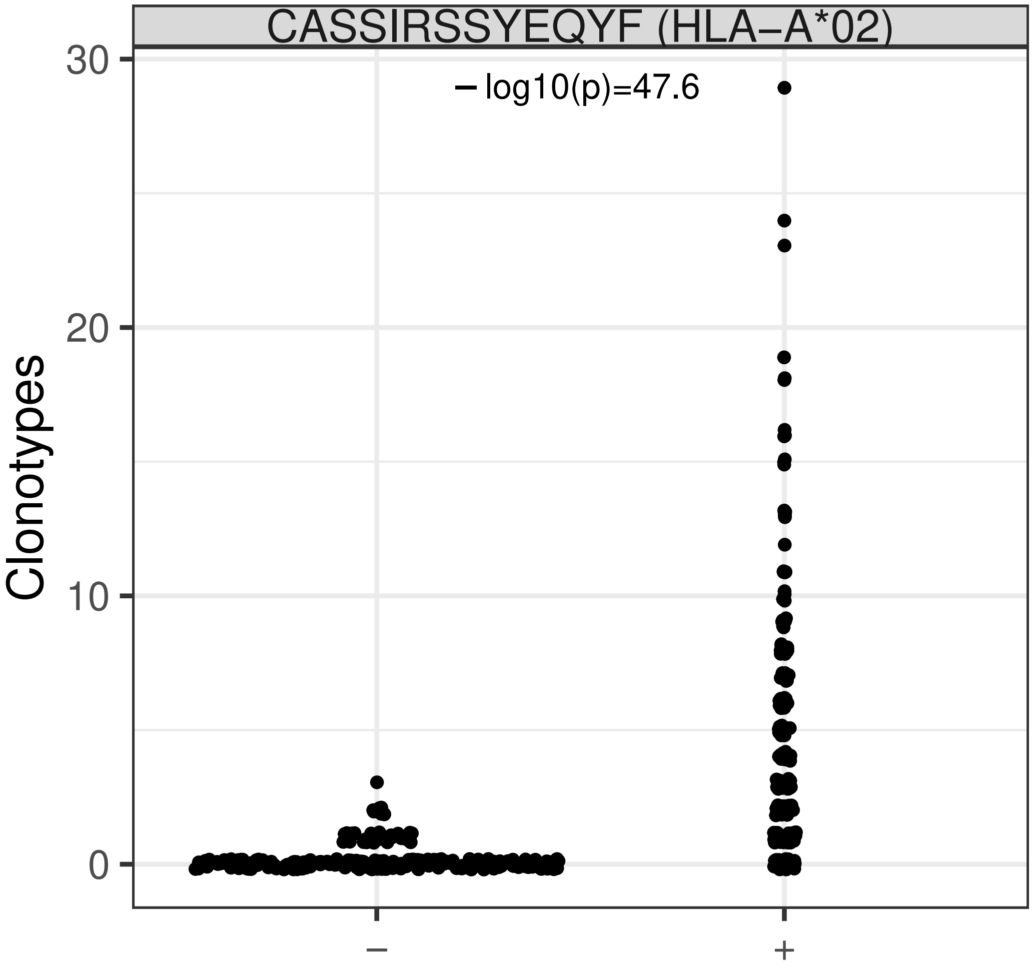


Supplementary Figure S1. Clonotype abundance of CDR3$\beta$ sequence CASSIRSSYEQYF in HLA-A*02-positive (n=241) and HLA-A*02-negative (n=267) individuals. Data from 508 TCR repertoires [Emerson et al., 2017] show individual clonotype counts (dots). Mann-Whitney U test p-value (-log10 scale) is labeled above the groups.

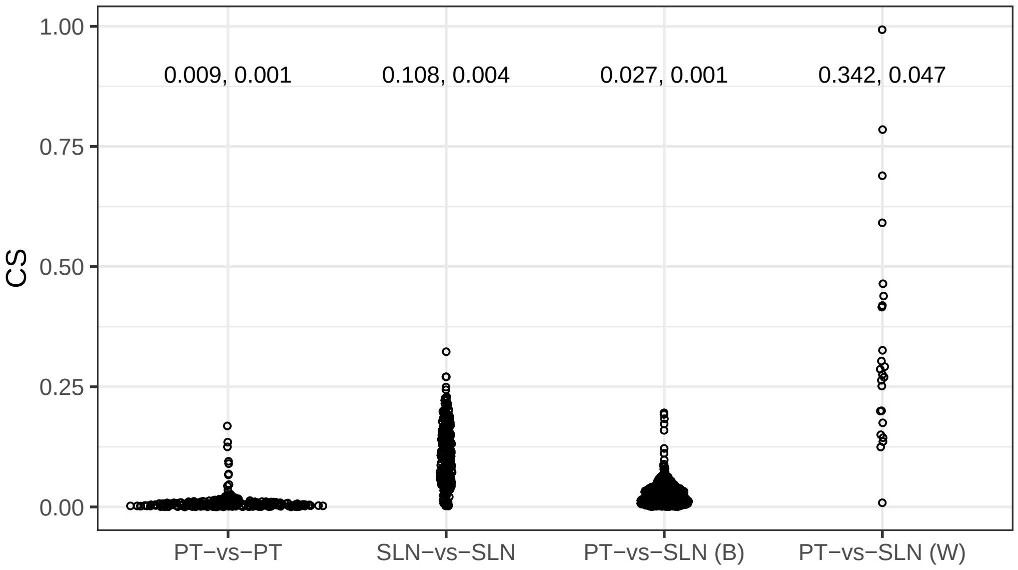


Supplementary Figure S2. Cosine similarity (CS) between CJ occupancies of pairs of TCR repertoires. Each dot represents the CS from a pairwise comparison between repertoires in one of four groups (x-axis): PT-vs-PT, both repertoires are from primary tumors (PT); SLN-vs-SLN, both repertoires are from sentinel lymph nodes (SLN); PT-vs-SLN (B), one repertoire is from a PT and the other from an SLN, with the two repertoires originating from different patients; and PT-vs-SLN (W), a within-patient comparison between PT and SLN repertoires. Labels indicate the mean $\pm$ standard error of the CS for each group.

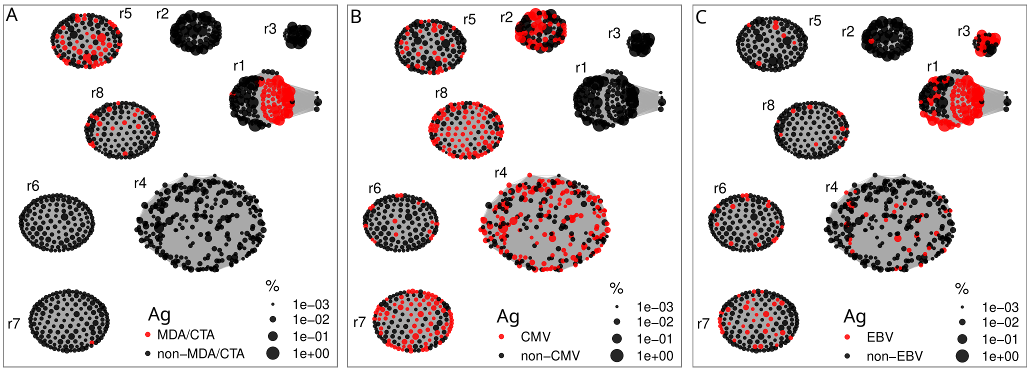


Supplementary Figure S3. Network of recurrent TCR communities (*r1-r8*). Nodes represent TCR clonotypes sized by relative clonal expansion. Gray edges connect clonotypes sharing similar CDR3$\beta$ sequences. Clonotypes specific and non-specific for MDA/CTA (panel A), CMV (panel B) and EBV (panel C) are shown as red and black, respectively.

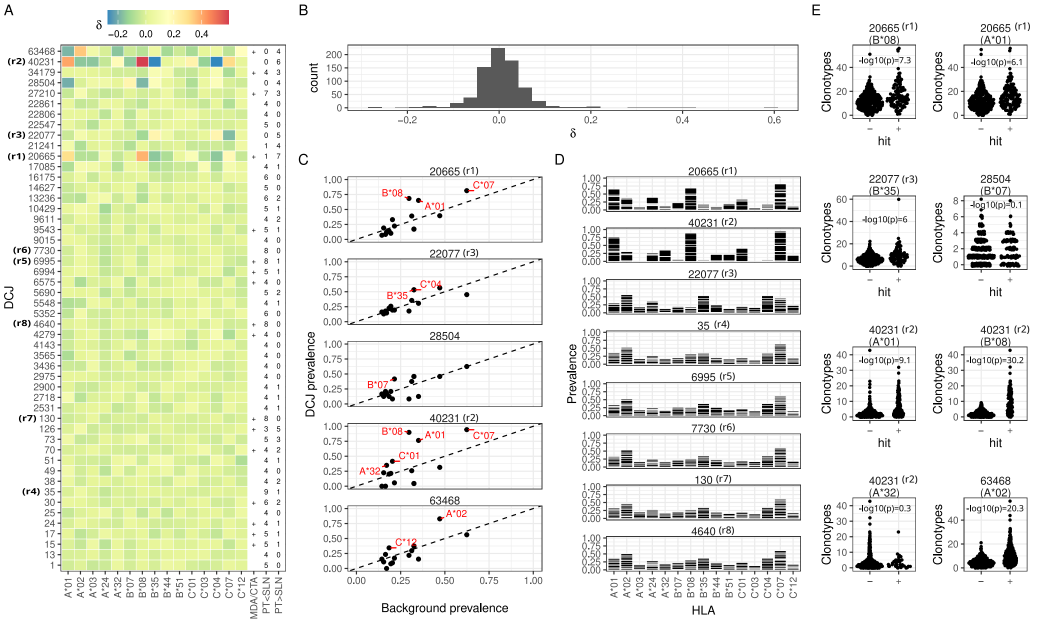


Supplementary Figure S4. HLA enrichment in 50 recurrent DCJ sets. (A) Enrichment heatmap ($\delta > 0$: enrichment; $\delta < 0$: depletion) for 15 HLA types (x-axis) across 42 DCJ sets (y-axis). Recurrent DCJ sets (*r1*–*r8*) are labeled; MDA/CTA-specific DCJ sets are marked with +. The number of primary tumors (PTs) and sentinel lymph nodes (SLNs) in which each DCJ set is differentially expanded is indicated in PT>SLN and PT<SLN, respectively. (B) Distribution of enrichment scores ($\delta$). (C) Expected (x-axis) vs. observed (y-axis) HLA prevalence in DCJ sets (panels); HLAs with $\delta > 0.15$ are labeled. (D) Relative clonotype abundance per patient (stacked bars) for each HLA type (x-axis) within each DCJ set (panels). (E) Abundance of clonotypes with CDR3$\beta$ sequences belonging to specific DCJ sets (panel titles) across TCR repertoires [Emerson et al., 2017] (dots) from individuals carrying (+) or not carrying (−) the HLAs indicated in the panel legend. Only HLA-A and HLA-B types are considered, as HLA-C data are only partially available in [Emerson et al., 2017]. Group comparisons were performed using the Mann-Whitney U test; $p$-values (-$\log_{10}$ scale) are shown above the groups.

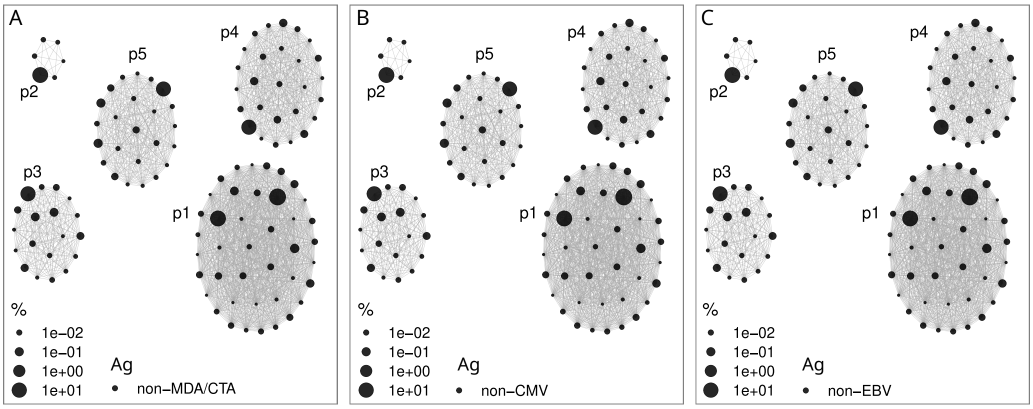


Supplementary Figure S5. Network of private communities (*p1-p5*): Nodes represent TCR clonotypes sized by relative clonal expansion. Gray edges connect clonotypes sharing similar CDR3$\beta$ sequences. Clonotypes specific and non-specific for MDA/CTA (panel A), CMV (panel B) and EBV (panel C) are shown as red and black, respectively.

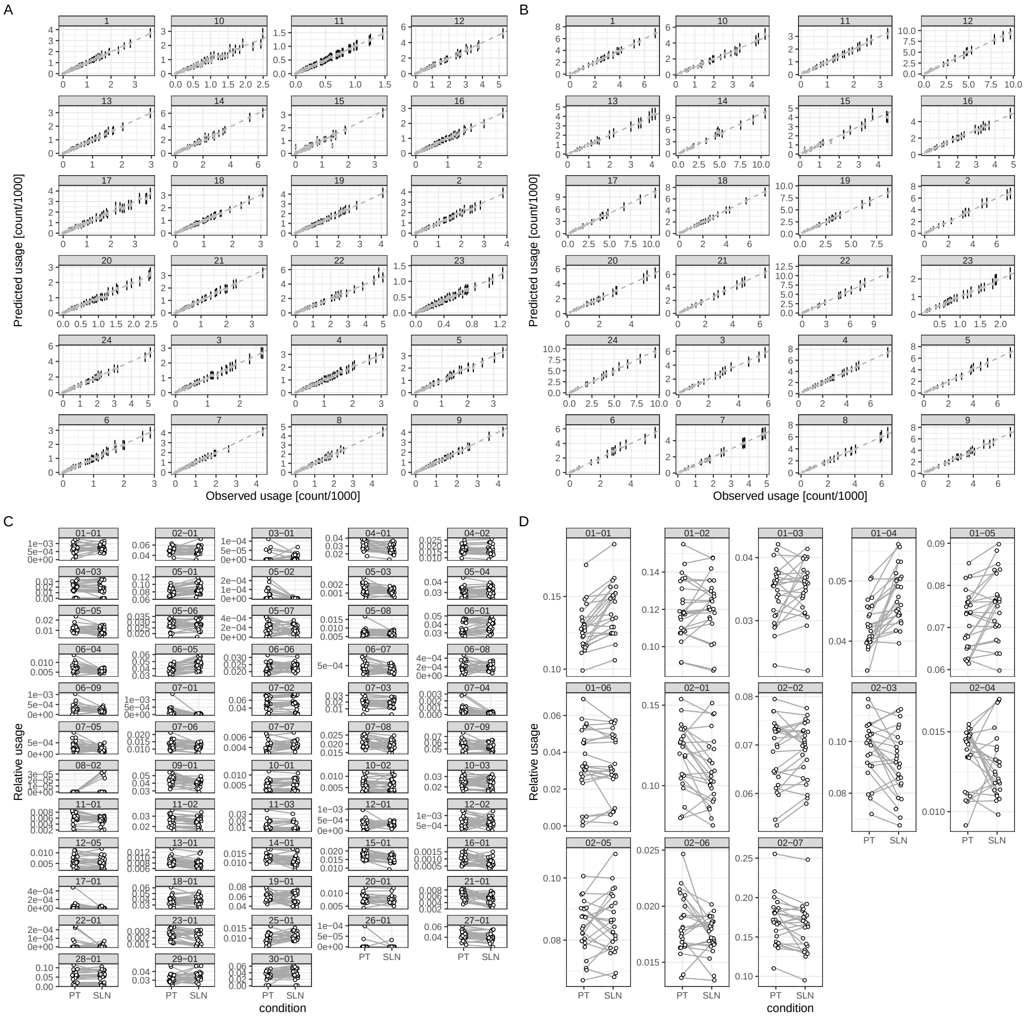


Supplementary Figure S6. Posterior predictive checks (PPCs). Observed (x-axis) and simulated mean usage (y-axis) and 95% HDI (error bars) for TRBV (A) and TRBJ (B) gene usage. Panels show TRBV (C) and TRBJ (D) gene usage (y-axis) in matching PTs and SLNs.

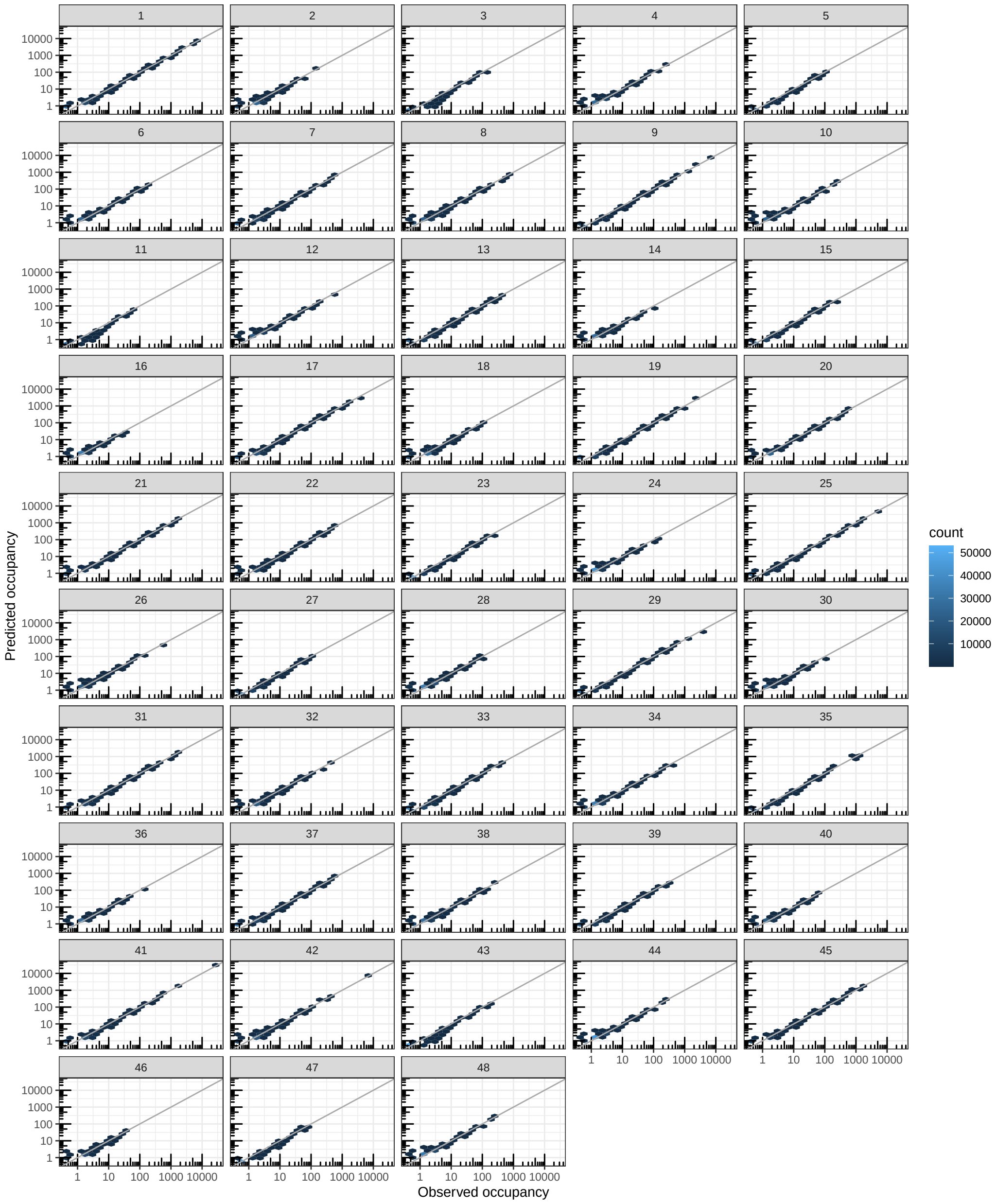


Supplementary Figure S7. Posterior predictive checks (PPCs) with *ClustIRR*. Observed (x-axis) versus simulated (y-axis) mean occupancy for 48 repertoires (panels). The color of the 2D hexagons indicates the density of communities in the observed–simulated occupancy space. The gray diagonal indicates perfect agreement between observed and simulated mean occupancy.
